# Characterizing the temporal dynamics of object recognition by deep neural networks : role of depth

**DOI:** 10.1101/178541

**Authors:** Kandan Ramakrishnan, Iris I.A. Groen, Arnold W.M. Smeulders, H. Steven Scholte, Sennay Ghebreab

## Abstract

Convolutional neural networks (CNNs) have recently emerged as promising models of human vision based on their ability to predict hemodynamic brain responses to visual stimuli measured with functional magnetic resonance imaging (fMRI). However, the degree to which CNNs can predict temporal dynamics of visual object recognition reflected in neural measures with millisecond precision is less understood. Additionally, while deeper CNNs with higher numbers of layers perform better on automated object recognition, it is unclear if this also results into better correlation to brain responses. Here, we examined 1) to what extent CNN layers predict visual evoked responses in the human brain over time and 2) whether deeper CNNs better model brain responses. Specifically, we tested how well CNN architectures with 7 (CNN-7) and 15 (CNN-15) layers predicted electro-encephalography (EEG) responses to several thousands of natural images. Our results show that both CNN architectures correspond to EEG responses in a hierarchical spatio-temporal manner, with lower layers explaining responses early in time at electrodes overlying early visual cortex, and higher layers explaining responses later in time at electrodes overlying lateral-occipital cortex. While the explained variance of neural responses by individual layers did not differ between CNN-7 and CNN-15, combining the representations across layers resulted in improved performance of CNN-15 compared to CNN-7, but only after 150 ms after stimulus-onset. This suggests that CNN representations reflect both early (feed-forward) and late (feedback) stages of visual processing. Overall, our results show that depth of CNNs indeed plays a role in explaining time-resolved EEG responses.

## 1 Introduction

The near-human performance of convolutional neural networks (CNNs) [19] on automated object recognition has led to a number of neuroimaging studies that investigated the correlation of CNNs to feedforward visual processing in the human brain. It has been shown that CNNs correlate much better to cortical representations measured with human neuroimaging than other computational models [18]. A similar correlation of CNNs was found to neural recordings from primate IT-cortex during core visual object recognition [2], [35]. Moreover, evidence suggests that CNNs map onto brain responses in a hierarchical manner, with lower CNN layers predicting responses in early visual cortex and high-level layers predicting responses in category-selective cortex [12], [4], [3]. While it is increasingly becoming clear that CNNs capture hierarchical representations in the human visual system, there are a number of open questions.

First, the impressive performance of CNNs in predicting brain responses has mostly been demonstrated for fMRI responses derived from slow fluctuations in blood flow across multiple brain regions in visual cortex [8]. However, object recognition is a fast process that is resolved within the initial hundreds of milliseconds of visual processing [32]. A cascade of visual processing stages gives rise to characteristic spatio-temporal dynamics shaped by both feed-forward and feedback processing [20] that can be measured with time-resolved magneto- and electro-encephalography (M/EEG). While CNN layers do not have a temporal dimension, it has been show that the CNN layers predict whole-brain decoding performance of MEG responses in the first few hundred milliseconds of visual processing in the human brain [4]. Interestingly, these results suggested that, just as in the spatial domain, CNN layers correspond hierarchically to temporally resolved responses, with lower layers predicting decoding performance early, and higher layers predicting performance later in time.

Second, CNNs have outperformed shallow computer vision models on automated object recognition datasets on ImageNet [26] and PASCAL VOC [5]. The performance on object recognition has further increased with deeper neural networks consisting of 15 layers (CNN-15) and 18 layers [30] compared to 7 layer CNN [19] on large image datasets. State-of-the-art CNNs are capable to discriminate between 1000 visual categories with error rates on par with human performance [13] and overlaps with human behavior [28]. The number of layers is a critical factor influencing the performance of the CNN architecture. For instance, while the CNN-7 of 5 convolutional layers achieves an error rate of 18% (top-5 error) on ImageNet, the CNN-15 with 13 convolutional layers achieves an error rate of 7.5% [30]. In both architectures, the layers consist of similar operations such as: filter convolution, non-linear activation, spatial pooling and normalization, with each network containing 2 fully connected layers.

**Figure 1:**
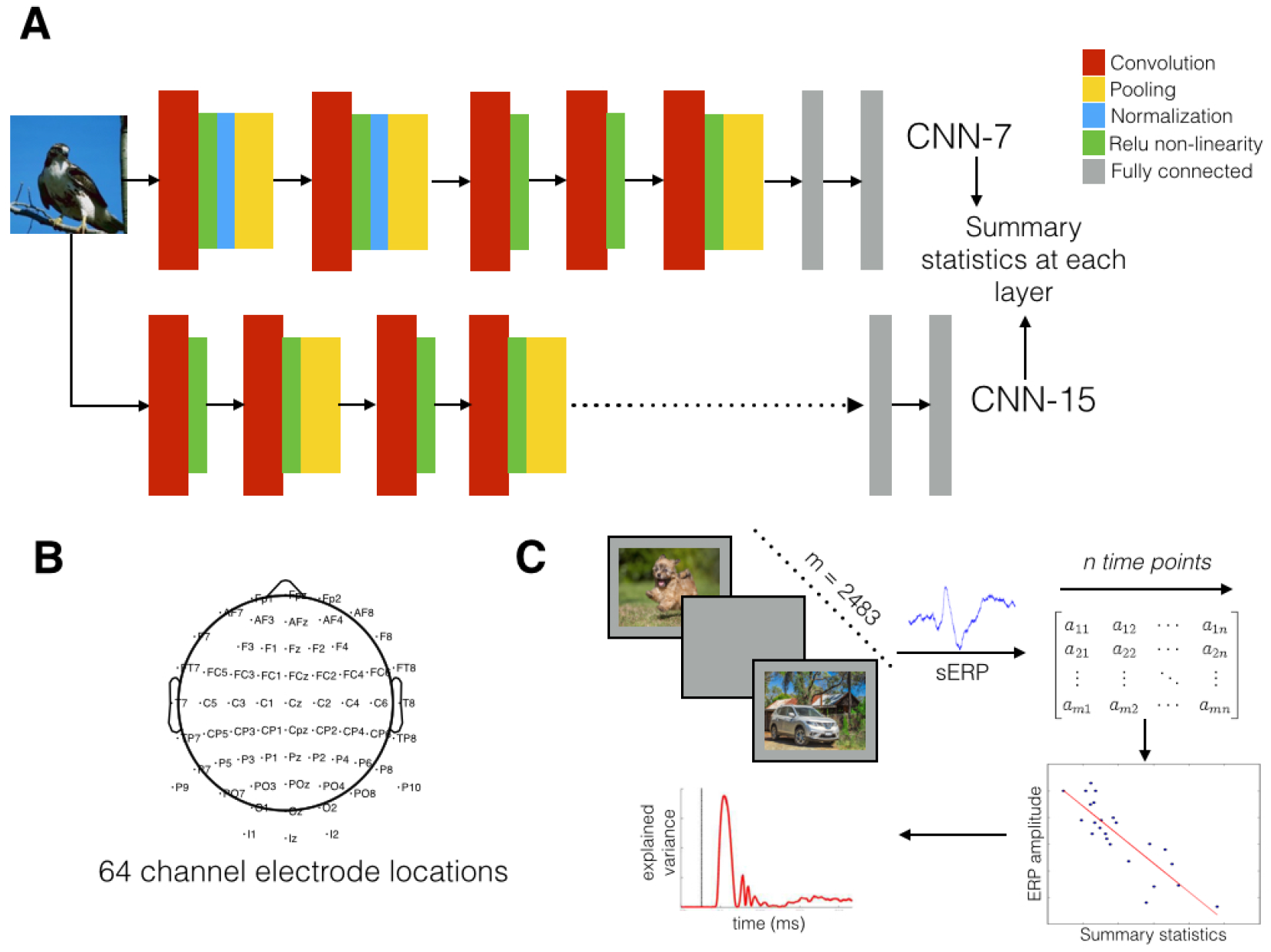
Models and experimental design : Fig. A. Architectures of a 7 (CNN-7) and 15 (CNN-15) layer CNN. CNN-7 consists of 5 convolutional layers and 2 fully-connected layers. The CNN-15 consists of 13 convolutional layers and 2 fully connected layers. Summary statistics (mean and mean/standard deviation) of the representations is computed at each CNN layer. B. The 64 channel EEG system with locations of the different channels. C. Single image ERPs at each channel, averaged over subjects, are regressed to the summary statistics of each layer from the CNNs by selection of 1800 images from the full set of 2483 images. The regression is permuted a 1000 times and this results in explained variance (r2) per layer for CNN-7 and CNN-15. We plot the average explained variance over the 1000 permutations per layer of CNN-7 and CNN-15.

The improved performance of CNN-15 compared to CNN-7 is commonly attributed to the additional non-linearity at the additional layers, which improves the discriminative power of the network. With increasing number of layers, the model contains one essential non-linear operator per layer, the rectified linear unit (abbreviated to relu). However, at the same time the CNN-15 has smaller filter sizes (analogous to receptive field sizes in the human brain) per layer. The effective field size of layer 1 in CNN-7 is the same over two layers combined in the CNN-15 architecture. In this way, a correspondence of effective receptive field sizes between different CNN architectures has been established [30]. Thus, the number of layers, number of non-linear relu’s and receptive field size are interrelated yet critical aspects of the CNN-architecture. The improved performance of deeper CNNs on automated object recognition raises the question whether these architectures also better model visual representations in the human brain.

In this article we examine whether the temporal dynamics of visual object processing in the human brain are better predicted by CNNs with increasing number of layers. To address this, we tested: 1) whether the hierarchy of CNN representations predict differences in visual evoked activity in a systematic manner over time and 2) if a 15 layer CNN model explains a higher variance of brain responses compared to a 7 layer CNN. To that end, we first measured event-related potentials (ERPs) responses of 20 individuals to a large set of natural images. The layers of different CNN models were regressed to each time point of the ERP responses to determine their ability to explain variance in evoked amplitude between individual images. We compared the different layers and different CNN models on the basis of explained variances obtained.

## 2 Materials and Methods

### 2.1 Subjects

Twenty-one participants (7 males, 22-33 years old, mean 25.6, = SD = 2.5) took part in the EEG experiment. All participants had normal or corrected-to-normal vision, provided written informed consent and received financial compensation. The ethics committee of the University of Amsterdam approved the experiment. Two subjects were excluded in preprocessing: one subject based on the presence of excessive alpha activity, and another because of a history of epilepsy.

### 2.2 Stimuli and Procedure

Participants viewed a large set of scene stimuli while performing go-no go object recognition tasks [9]. A stimulus set of 6800 color images (bit depth 24, JPG format, 640 x 480 pixels) was composed from several existing online databases. The set included images from a previous fMRI study on scene categorization [34], as well as images from various datasets used in computer vision: the INRIA holiday database [16], the GRAZ dataset [22], ImageNet [26], and the McGill Calibrated Color Image Database [21]. These different sources assured maximal variability of the stimulus set: it contained a wide variety of indoor and outdoor scenes, landscapes, forests, cities, villages, roads, images with and without animals, objects, and people. For the purpose of the current study, we analyzed only the images that are not contained in the ImageNet dataset, resulting in 2483 number of images. This was to avoid overlap in training deep neural networks and the stimuli presented to the participants.

Stimuli were presented on a 19-inch Ilyama CRT-monitor (1024×768 pixels, frame rate 60 Hz). Participants were seated 90 cm from the monitor such that stimuli subtended 14×10 deg of visual angle. On each trial, one image was randomly selected and presented in the center of the screen on a grey background for 100 ms, on average every 1500 ms (range 1000 – 2000 ms). In different task blocks, participants searched for either animals or vehicles at four levels of categorization: basic detection (animal/no-animal and vehicle/no-vehicle), superordinate (animal/vehicle and vehicle/animal), basic-level (cat/other animals and bicycle/other vehicles) or subordinate categorization (Persian cat/other cats and mountain bike/other bicycles). Within each task, subjects performed a total of 800 trials (400 target, 400 non-target images). In addition, subjects performed an intact vs. scrambled scene task on a subset of 400 scene images and their Fourier phase-scrambled counterparts (only the intact scenes were analyzed). For each participant, data were obtained across three separate recording sessions that were conducted on different days. Per recording session, each participant performed each of the 7 different tasks for a particular subset of the stimuli. Task orders and stimulus subsets were counterbalanced across participants and recording sessions. Participants were instructed to respond as quickly and accurate as possible, and indicated their responses with their right hand using a custom-made button box that was taped to the chair armrest. Prior to the start of each task block, participants performed 20 practice trials on images that were not included in the main experiment. Each task block was interspersed with a short break allowing subjects to rest. Stimuli were presented using the software package Presentation (www.neurobs.com).

### 2.3 EEG acquisition and preprocessing

EEG Recordings were made with a Biosemi 64-channel Active Two EEG system (Biosemi Instrumentation BV, Amsterdam, NL, www.biosemi.com). Recording set-up and preprocessing were identical to the procedures described in [11], [10]. We used caps with an extended 10-20 layout, modified with two additional occipital electrodes (I1 and I2, while removing electrodes F5 and F6). During recording, a CMS/DRL feedback loop was used as an active ground, followed by offline referencing to electrodes placed on the earlobes. The Biosemi hardware is completely DC-coupled, so no high-pass filter is applied during recording of the raw data. A Bessel low-pass filter was applied starting at 1/5th of the sample rate. Eye movements were monitored with a horizontal electrooculogram (hEOG) placed lateral to both eyes and a vertical electro-oculogram (vEOG) positioned above and below the left eye, aligned with the pupil location when the participants looked straight ahead. Data was sampled at 256 Hz. Pre-processing occurred in Brain Vision Analyzer and included a high-pass filter at 0.1 Hz (12 dB/octave); a low-pass filter at 30 Hz (24 dB/octave); two notch filters at 50 and 60 Hz; automatic removal of deflections > 300 mV; epoch segmentation in -100 ms to 500 ms from stimulus onset; ocular correction using the EOG electrodes [7]; baseline correction between -100 ms and 0 ms; automated artifact rejection and conversion to Current Source Density responses [23]. No trial or electrode averaging was performed: preprocessing thus resulted in a single-trial EEG response specific to each subject, electrode and individual image presentation. Prior to regression analysis, responses were averaged across participants, resulting in a single event-related potential (ERP) specific to each individual image.

### 2.4 CNN representations and summary statistics

Our analysis consisted of two stages. In the first stage, we used two different pre-trained CNN architectures: the 7-layer CNN-architecture (CNN-7) [19] and the 15-layer CNN-architecture, CNN-15 [30](Figure 1A). The CNNs used in this study were pre-trained on the ImageNet dataset with the same hyper-parameters as described in the MatConvNet toolbox http://www.vlfeat.org/matconvnet/pretrained/.

In the second stage, we summarized the representation of each CNN layer by two parameters – the mean and mean normalized by standard deviation (Figure 1B). These summary statistics have been previously found to constitute a biologically plausible model of population receptive field outputs [27] and have been used successfully for-natural image identification based on EEG-responses [6] and to describe the population activity captured by individual EEG electrodes[10]. Additionally, the use of summary statistics allows us to better handle the high dimensionality of the CNN feature representations, and to equate the number of parameters extracted for each CNN layer.

### 2.5 Regression of CNN layers on single image ERPs

To test whether differences between evoked neural responses could be predicted by the summary parameters of the CNNs, we conducted regression analyses on the single-image ERPs (Fig. 1C). The preprocessed ERPs were used in Matlab, where we conducted linear regression analyses of ERP amplitude on the CNN layer summary parameters using the Statistics Toolbox. At each channel and each time-point, two summary parameters for each CNN layer were entered together as linear regressors on single-image ERP amplitude. This analysis results in a measure of model fit (adjusted r2, corrected for predictor dimension) over time (each sample of the ERP) and space (each electrode). The fit of the regression model was statistically evaluated by permutation tests: we randomly selected 1800 out 2483 images and repeated the regression analysis a 1000 times. We averaged the r2 over all permutations to represent the correlation between CNN layers and the ERP amplitude.

### 2.6 Statistical testing

For significance testing of the explained variance, the permutation analysis (1,000 times) was used to calculate the standard deviation of the sampled bootstrap distribution. To correct for multiple comparisons across permutations, time-points, channels and layers, we used the Bonferroni measure, resulting in an adjusted alpha = 1e-10. To compare the results directly (comparing layers within each model), we used the Akaike information criterion (AIC), which measures the information contained in each set of predictors (summary statistics of each CNN layer). Specifically, we transformed the residual sum of squares (RSS) of the regression analysis based on each set of statistics into AIC-values using AIC = n*log(RSS/n)+2k where n = number of images and k is the number of predictors. AIC can be used for model selection given a set of candidate models of the same data, where the preferred model has minimum AIC-value [1]. Thus in comparing the different CNN layers, the CNN layer with the lowest AIC value has the best fit to evoked activity. To compare the two models directly, we use a paired t-test using the permuted values at each time point.

### 2.7 Regression of combined CNN layers on single image ERPs

To test the performance of each CNN model in their entirety (rather than layer-by-layer), we repeated the regression analysis using the combined summary statistics of all layers in each CNN architecture. This resulted in two different regressors for each of the two CNNs: the CNN-7 regressor was of dimension 14 (2 parameters for each layer) and the CNN-15 was of dimension 30. Similar to the separate, individual CNN layer regression analyses, this analysis resulted in the measure of model fit (adjusted r2, which is corrected for differences in the predictor dimension of the two architectures) over time and space for the average subject response. As before, statistical evaluation was done by permutation testing.

### 2.8 Comparison of CNN models

To better understand the correspondence between the two CNN architectures, we correlated the summary statistics of the layers across architectures (architecture correspondence). We also correlated the time series of the explained variance (r2) of all the electrodes (converted to a vector consisting of time points of all electrodes) from each layer of different CNNs (neural correspondence). Specifically, the vector of explained variances at each layer of CNN-7 was correlated to each layer of the CNN-15 (Pearson correlation).

## 3 Results

### 3.1 Maximal correlation of individual CNN layers

We first examined the entire time-courses of explained variance of the ERP responses by the CNN architectures. Figure 2A shows the time point of maximal explained variance for the different CNN-7 and CNN-15 layers across all channels. Figure 2B displays the explained variance by both CNN architectures across the entire scalp. We observed that for CNN-7, layers 1-5, the maximum explained variance was found between 110 ms and 120 ms. For CNN-15, layers 1-10 of CNN-15 reached maximal variance between 110 and 120 ms. From Figure 2B we observed that the maximum explained variance early in time was localized to the electrodes overlaying the occipital cortex for CNN-7 (r2 = 0.43 for layer 1) and CNN-15 (r2 = 0.41 for layer 5).

However, for the higher CNN layers (layer 6 for CNN-7 and layer 12-15 for CNN-15), the maximum explained variance was found later in time between 160ms and 210ms. Furthermore, from the scalp visualization plot in Figure 2B the maximal explained variance later in time is observed at channels overlaying lateral-occipital cortex.

**Figure 2:**
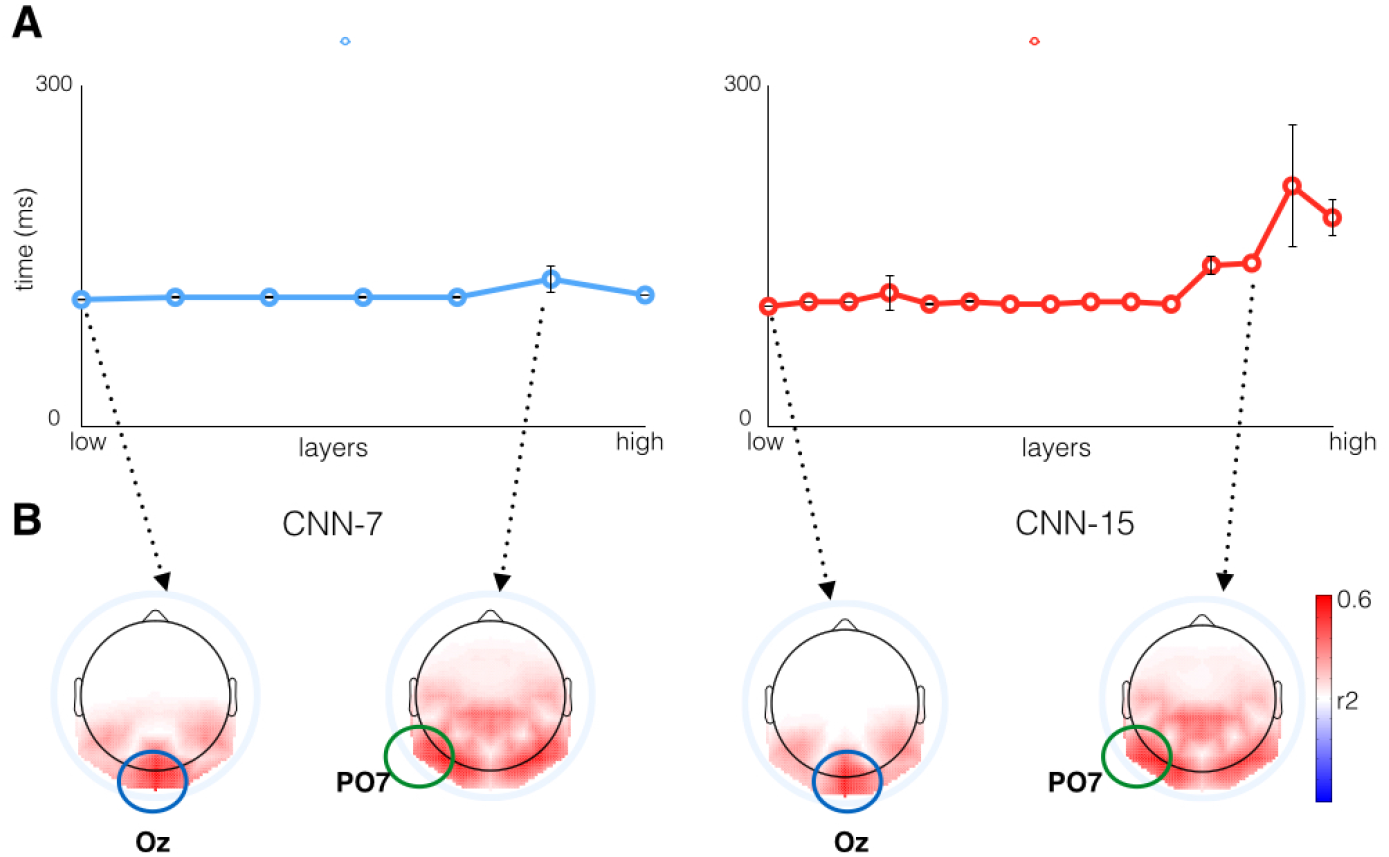
Peak explained variance of CNN layers. A) The time point of peak explained variance by the CNN-7 layers over the entire scalp. Lower CNN layers had maximum explained variance early in time and higher CNN layers later in time. We observed from the scalp visualization that the highest explained variance early in time was found at the occipital channel Oz. Similarly, highest variance later in time was observed at the peri-occipital channel PO7. B) Visualization of the time point of maximum explained variance by the CNN-15 layers over the entire scalp. Similar to CNN-7, lower CNN layers of CNN-15 reached maximum explained variance early in time at Oz and higher CNN layers later in time at PO7.

Our results show that the hierarchy of CNN representations correlate to the temporal hierarchy of visual processing and generalizes across different architectures. Low- and high-level CNN representations explain variance in visual evoked responses across multiple time-points. Moreover, while ERP responses generally have poor spatial resolution, we observe some spatial specificity in the pattern of results, with higher layers corresponding best to response differences at peri-occipital electrodes overlaying lateral visual areas, whereas lower layers corresponded to response differences at occipital electrodes overlaying early visual regions (SI Video 1).

### 3.2 Time course of CNN layer correlation to ERP

To examine the spatial specificity of our results, we focused our remaining analyses on two specific electrodes: the occipital channel Oz and peri-occipital channel PO7, as these electrodes exhibited strongest correlation with early and late CNN layers. Note that this is in line with previous studies which indicated sensitivity of summary statistic models to both early occipital (Oz) and late peri-occipital (PO7, PO8) channels [11], [27].

#### 3.1 Occipital channel

CNN-7 explained a substantial amount of variance in ERP responses to individual images as shown in Figure 3A. The maximum explained variance was *r*2 = 0.46 at 117 ms for the occipital channel Oz, specifically by CNN layer 1. Explained variance for all layers reached a maximum between 110 and 120 ms after stimulus onset; maximal values for the lower CNN layers ranged between r2 = 0.40 – 0.45 (*p* < 1*e* – 10 for t > 100*ms*, Bonferroni-corrected for multiple comparisons). The explained variance of EEG responses by CNN layers 6 and 7 were lower, r2 = 0.20 – 0.25 (*p* < 1*e* – 10 for *t* > 100*ms*, Bonferroni-corrected for multiple comparisons). The AIC values for the different layers are displayed in Figure 3B. CNN layer 1 had the lowest AIC values for 100 ms to 150 ms, while higher CNN layers had higher AIC values. This result indicates that the lower CNN layers best explained responses at the occipital channel.

**Figure 3:**
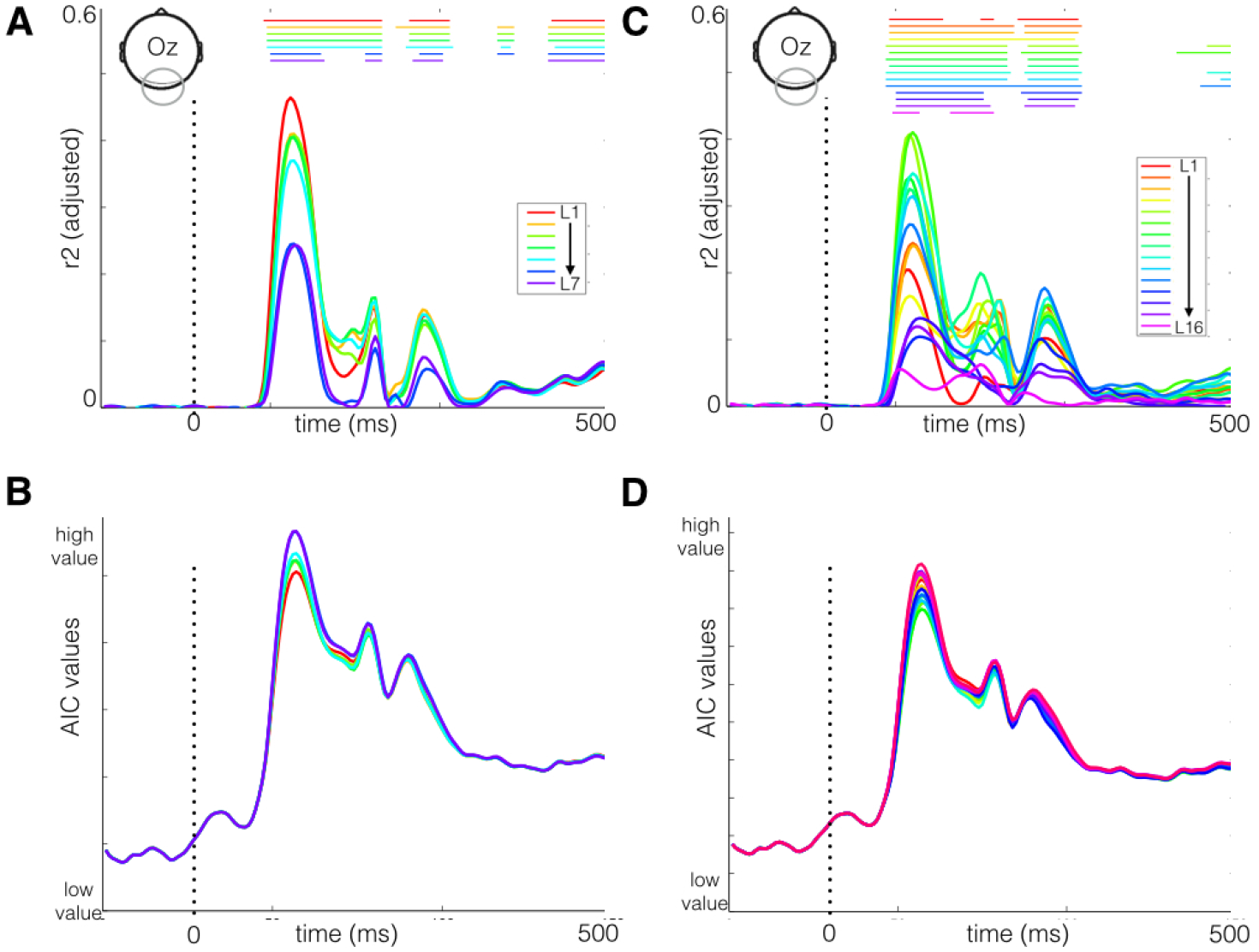
EEG regression results at electrode Oz. A) Explained variance at the occipital channel for the different layers of the CNN-7 architecture. Significant p-values (*p* < 1*e* – 10 from the permutation analysis) corresponding to each layer are displayed at the top of the graph. All layers correlated with evoked activity starting from 100 ms after stimulus onset. B) AIC values computed from the residuals of each of the CNN layers, showing that layer 1 provided the best fit to the data (low AIC value). C-D) Same as A-B, but for CNN-15. Similarly to CNN-7, lower layers of CNN-15 were maximally correlated to the ERP responses between 100-300 ms, with layer 5 resulting in the best fit at Oz.

In comparison, CNN-15 also explained a substantial amount of variance as shown in Figure 3C. Similar to CNN-7, the explained variance of ERP responses by CNN-15 in the occipital channel Oz was maximal between 110 and 120 ms after stimulus onset. All the lower layers were significantly correlated (*p* < 1*e* – 10) between 100 – 200ms. Specifically, layer 5 and 6 from CNN-15 had maximum explained variance r2 = 0.40 – 0.43 (*p* < 1 *e* – 10, Bonferroni-corrected for multiple comparisons) at the occipital channel. The AIC values in Figure 3D show that layer 5 had the lowest value between 100-150 ms, similar to the results for CNN-7.

Together, these results demonstrate that lower layers in both CNN-7 and CNN-15 best modeled evoked activity at the occipital channel with the explained variance reaching a peak between 110 and 120 ms. Moreover, at this channel and time window the correspondence between the evoked activity and the CNN decreased with increasing layers, revealing an (inverse) hierarchical mapping of CNN depth to brain responses early in time at activity recorded on electrodes overlaying low-level visual areas.

#### 3.2 Peri-occipital channel

Next, we considered the results for the peri-occipital electrode, which showed a different pattern of results relative to the occipital electrode (Figure 4). Specifically, higher CNN-7 layer 6 had maximum explained variance later in time (*t* > 160*ms* after stimulus onset). Out of all layers, layers 5 and 6 yielded the highest explained variance which ranged between r2 = 0.20 – 0.25 (*p* < 1*e* – 10, Bonferroni-corrected for multiple comparisons) between 160 and 170 ms after stimulus onset (Figure 4A). Critically, lower CNN layers showed no significant correlation with the ERP responses, demonstrating selective sensitivity to higher CNN layers at this electrode. Moreover – in contrast to the occipital channel – the AIC values indicated that higher CNN layers had the lowest values (Figure 4B). These results suggest that at higher-level lateral-occipital electrodes, higher CNN layers provide the best fit to ERP responses specifically later in time.

**Figure 4:**
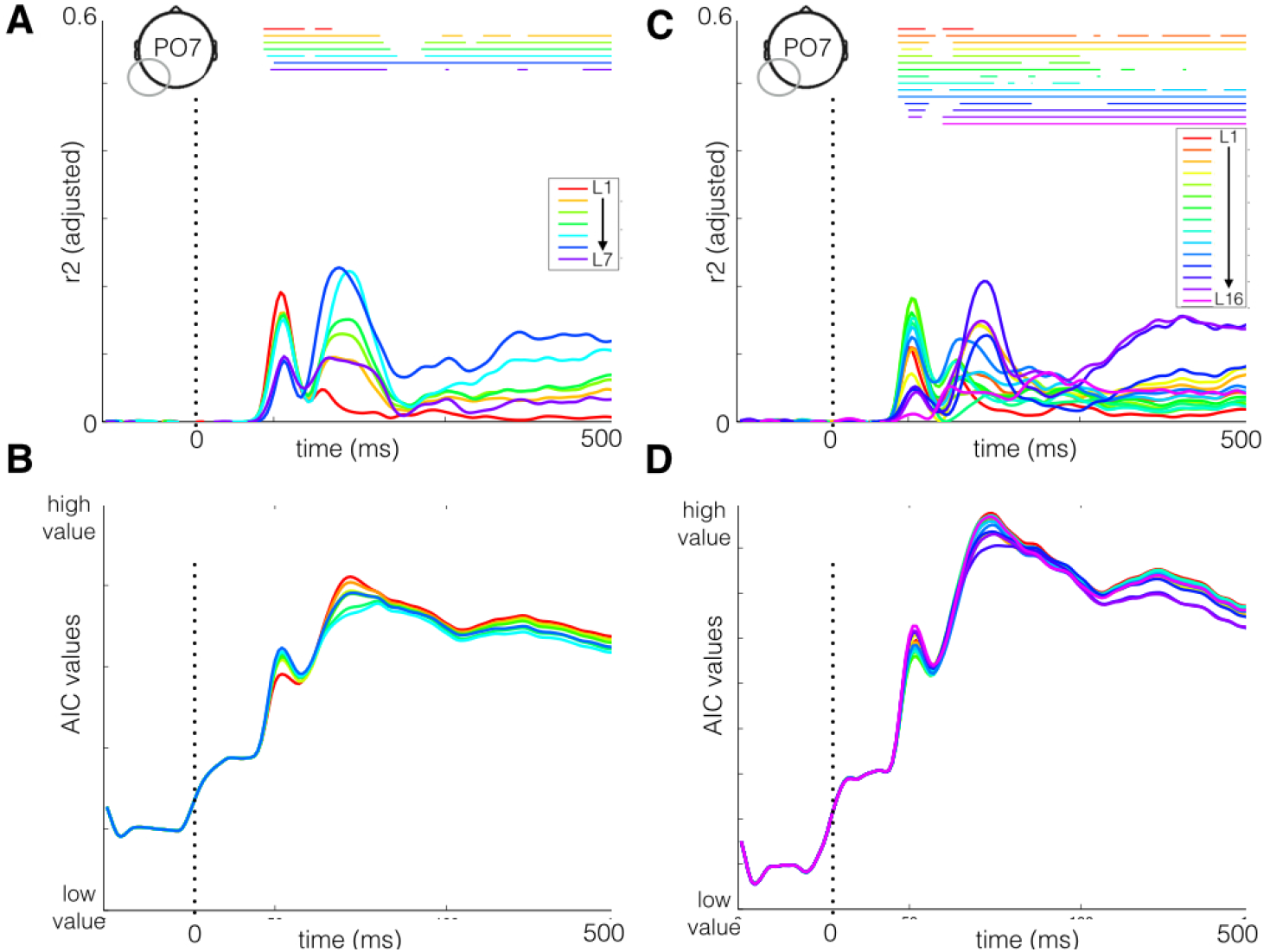
EEG regression results at electrode PO7. A) Explained variance at the peri-occipital channel for the CNN-7 architecture. Lower CNN layer 1 and 2 did not correlate to ERP responses, while layer 3-6 were significant after 150 ms (*p* < 1*e* – 10). B) AIC values for each of the CNN layers. Layer 6 provided the best fit to the ERP responses (low AIC value). C-D) Same as A-B but for CNN-15, showing that layer 14 provides the best fit at the peri-occipital channel.

The results for CNN-15 were similar to those for CNN-7: explained variance reached a maximum between 165 and 180 ms for higher CNN layers (Figure 4C). The maximum explained variance of r2 = 0.21 (*p* < 1*e* – 10, Bonferroni-corrected for multiple comparisons) was found for layer 14. Interestingly, the maximum explained variance of ERP responses by CNN-15 is slightly lower than those obtained for CNN-7. Furthermore, the lower CNN layers were significant (*p* < 1*e* – 10) only early in time (between 100-140ms) with lower explained variance up to r2 = 0.18. Thus, similar to CNN-7 the higher layers of CNN-15 best model ERP responses later in time.

Overall, our results clearly show that early in time, ERP responses were best explained by lower CNN layers (with maximum correspondence in the occipital channel). In contrast, later in time, ERP responses were best explained by the higher CNN layers (with maximum correspondence at the peri-occipital channel). While the exact distribution of explained variance time course across layers for CNN-7 and CNN-15 showed subtle differences, the gradient of the layers (lower layers to higher in Oz and higher layers to lower in PO7) was similar for both CNN architectures. This suggests that the correlation of CNNs to the temporal hierarchy of visual representations does not fundamentally change with the number of layers. Surprisingly, we did not observe differences in the maximum explained variance between CNN-7 and CNN-15. This suggests that while CNN-15 is more complex than CNN-7 by virtue of more number of layers in the network, the addition of individual layers seems to have little added value for the temporal correlation. In the next section, we further examine potential differences between the CNN architectures by combining the representations at different layers.

### 3.3 Combined CNN layers

To investigate to what extent the total CNN architectures capture ERP responses, we concatenated summary statistics of all CNN layers and used these concatenated vectors as regressors. We then obtained the explained variance as before, while correcting for the difference in feature dimensions by using the adjusted r2 values.

Figure 5A shows the explained variances of ERP responses at the occipital electrode. Before 150 ms, combining layers did not yield a significant change in explained variance by CNN-7 and CNN-15. This is further seen in the AIC values, in which there is no difference between the architectures (Figure 4B). The explained variance reached a maximum of r2 =0.48 (*p* < 1*e* – 10, Bonferroni-corrected for multiple comparisons), between 100 and 110 ms after stimulus onset. This was observed for both CNN-7 and CNN-15. However, differences between the two CNN architectures emerged later in time (after 150 ms), with the combination of CNN-15 layers explaining more variance of ERP responses. This is also observed in the AIC values in Figure 5B, which show that the deeper CNN provided a better fit to ERP responses compared to CNN-7 between 150 and 400 ms. After 150 ms, the maximum explained variance by CNN-15 is r2 = 0.32 which is significantly higher than the maximal explained variance by CNN-7 (paired t-test, *t*(99) = 67.75, *p* < 0.05, *ci* = 8 – 10%). Thus, differences between CNN-7 and CNN-15 were observed later in time in the occipital channel.

**Figure 5:**
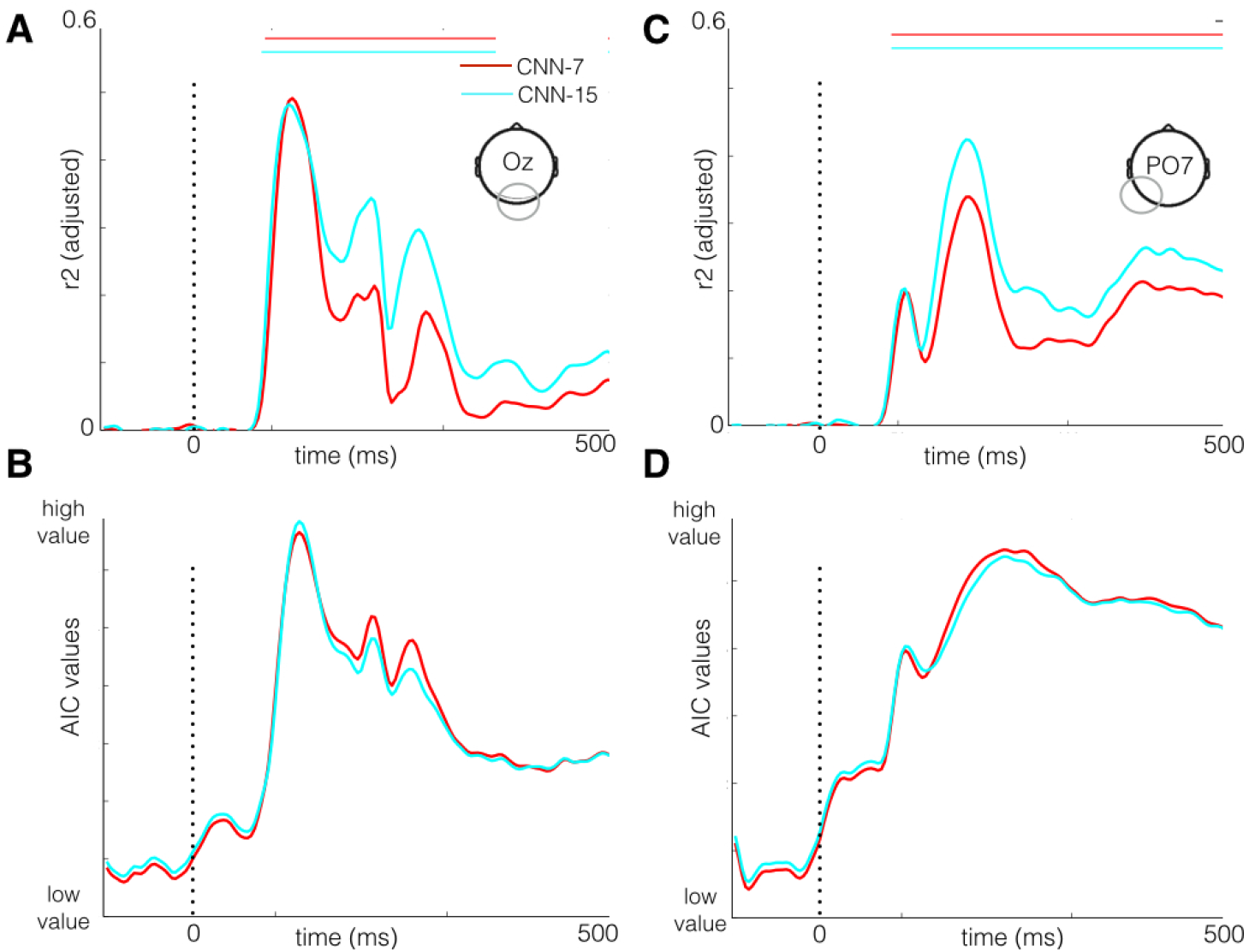
Comparison of CNN-7 and CNN-15 total architectures. Total architectures were obtained by combining the features from the different layers in a single regression model for each CNN. A) Explained variance at the occipital channel for the CNN-7 and 15 architecture (*p* < 1*e* – 10). B) AIC values for CNN-7 and CNN-15, which shows that the CNN-15 provides a better fit to the ERP values. C-D) Same as A-B but for the peri-occipital channel, showing that CNN-15 provides the better fit compared to CNN-7.The figures show that while there was no difference between the architectures early in time, deeper CNNs provided a better fit later in time.

A similar pattern of results was obtained for the peri-occipital channel PO7 (Figure 5C). For CNN-7 we observe a significant increase in explained variance reaching a maximum of r2 = 0.33 (*p* < 1*e* – 10, Bonferroni-corrected for multiple comparisons) as seen in Figure 5B. The maximum explained variance of the individual layers of CNN-7 reached r2 = 0.21 at *t* = 168ms. This is also observed for the CNN-15 total model, with a significant explained variance of r2 = 0.41 (*p* < 1*e* – 10, Bonferroni-corrected for feature dimension) compared to the variance by individual layers of r2 = 0.20 (*p* < 0.05). Importantly, the maximum explained variance of the CNN-15 is significantly higher than CNN-7 (paired t-test, *t*(99) = 62.58, *p* < 0.05, *ci* = 8 – 10%). From the AIC values, as shown in Figure 5D, the CNN-15 provides a better fit to the ERP responses compared to CNN-7 after 150 ms. It is worth noting that the increased explained variance is not due to the increase in number of parameters. This can be observed from the baseline before stimulus onset, which remains zero.

In sum, we observed that the combination of CNN layers gave rise to significant differences between the CNN architectures, with deeper CNNs providing a better fit to model brain responses. However, this difference was observed only later in time by combining lower and higher CNN layers in a single model.

### 3.4 Comparison of CNN architectures

To better understand what drives the observed difference in performance by the combination of CNN representations, we next performed an analysis that directly compared the two CNNs. We correlated the representations contained in each layer (Figure 6A) as well as their entire spatio-temporal profile of correspondence with evoked activity (Figure 6B; see Methods).

**Figure 6:**
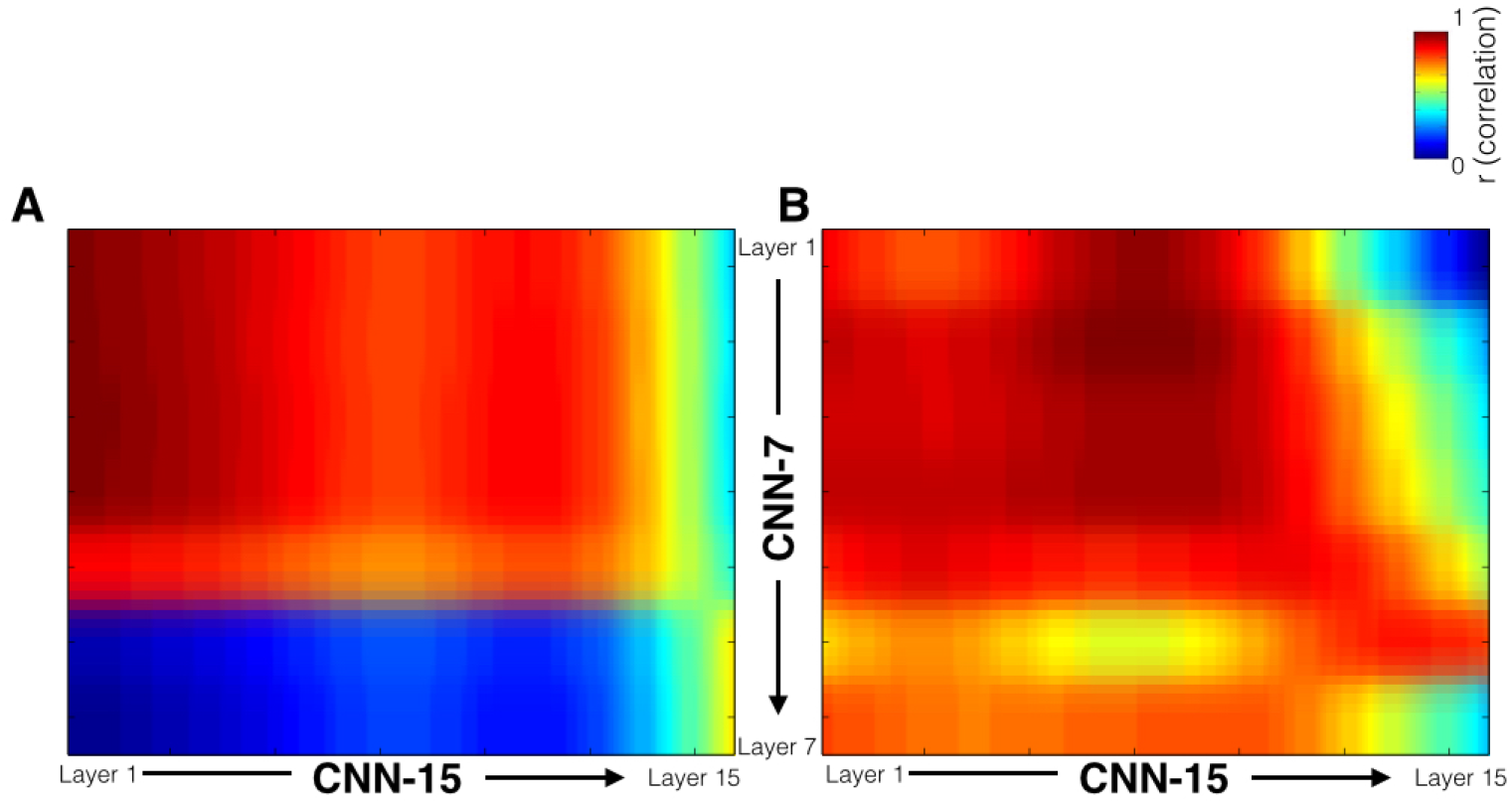
Visualization of the correspondence between the CNN-7 layers with CNN-15 layers. A) Architectural correspondence: Visualization of the correlation (*p* < 0.0005) between representations of each CNN-7 layer to each of the CNN-15 layer. The lower layers are highly correlated to each other across CNN-7 and CNN-15, compared to the correlation between higher layers. B) Neural correspondence: The explained variance by each layer is concatenated across all channels into a single vector, which is then correlated to the other layers. A non-linear relationship is observed for the correlation between the lower layers of CNN-7 and CNN-15.

In Figure 6A, we observe a hierarchical mapping of the representations contained in each layer across the two CNN architectures. The lower layers across architectures correlated most strongly (*p* < 0.0005, Bonferroni-corrected for multiple comparisons). These correlations decreased for higher layers across CNN architectures. This suggests that lower layer representations are highly similar across architectures, while higher layers are less similar. This is not surprising, while the lower layers learn similar features (gabor-like) across the architectures, the higher layers of CNN-15 are more discriminative compared to CNN-7. Further, we observe that the convolutional layers and fully connected layers correlate correspondingly across the two architectures.

While the neural correspondence (Figure 6B) also shows a hierarchical mapping, we also observe some differences in contrast to Figure 6A. As before, the highest layers of CNN-15 correlated to the higher CNN-7 layers (*p* < 0.0005), suggesting they captured similar stages of the neural processing. However, the spatiotemporal profile of explained variance by layer 7 of CNN-7 correlated most strongly with lower-and intermediate layers of CNN-15 (*p* < 0.0005, Bonferroni-corrected for multiple comparisons). Additionally, intermediate layers of CNN-15 correlated more strongly to lower layers of CNN-7 (*p* < 0.0005, Bonferroni-corrected for multiple comparisons). This suggests that the expansion of layers in CNN-15 relative to CNN-7 is reflected in low-level layers based on neural data : that is, the same neural response variance captured by low-to-intermediate layers in the CNN-15 is explained by the lowest CNN-7 layers.

In sum, together these results show that while CNN-7 and CNN-15 have similar architectural mapping due to design (i.e. lower CNN layers correlate with other to lower layers, and higher layers with higher layers), this relationship is not directly reflected in the correlation of CNN layers to neural responses, in which we observe a shift from low-to-intermediate layers in terms of the neural information processing captured by the CNN.

## Conclusion

The success of Convolutional Neural Networks (CNN) in computer vision has led to a number of demonstrations of a correspondence between CNNs and the brain [18, 2, 35, 12]. Most of these studies have focused on the correlation of CNN layers to neuromaging data using fMRI. However, object recognition is also reflected in time-resolved neural responses. A recent study [4] demonstrated a hierarchical mapping in the temporal domain of CNN layers to whole-brain decoding of visual representations of objects. Our study provides additional quantitative evidence that CNN-models are able to predict spatio-temporal dynamics of visual processing in humans. We confirm the previous finding from [4] that lower CNN-layers correlate to brain responses early in time while the higher CNN-layers correlate to responses later in time. Moreover, we find a subtle spatial shift in terms of these results, with lower layers primarily explaining ERP amplitude differences at occipital channels overlying early visual regions, while higher layers best explain differences at channels overlying higher-level lateral-occipital regions involved in object representations [17]. Thus, similar to the spatial mapping of hierarchical CNN representations to fMRI data from multiple studies, we observe a temporal correspondence of CNN representations to evoked EEG responses.

Our results further demonstrate generalization to deeper CNNs: we find highly similar results with CNNs containing 7 layers or 15 layers. There are no clear differences in maximal explained variance between the different CNNs when comparing individual layers. However, when we combined the different layers of the CNN in a single regression model, CNN-15 did result in a better fit of neural activity compared to CNN-7, but selectively later in time (i.e., beyond 150 ms). In this aspect, the deeper model is more powerful: they capture more information than the shallower CNN. Direct comparison of the representations and spatio-temporal mapping of CNNs to the neural responses suggested that this additional information was mostly reflected as more extensive representations of low-to-intermediate features in CNN-15. Overall, we demonstrate that CNN with larger numbers of layers are better models to explain temporal visual processing than shallow CNN-models at later, but not early stages of visual processing.

What may account for the temporal hierarchy we observed in the CNN correspondence to ERP responses? We speculate that the close correspondence of CNN-layers to the temporal dynamics of human object recognition might be attributed to the hierarchical nature of representations in the model [36]. The stacking of linear-nonlinear operations in the CNN closely resembles the simple-complex cell model as proposed as Hubel and Wiesel [15]. In the first two convolutional layers of the CNN, oriented gradient features similar to the receptive fields in the V1 and the V2-regions of the brain are learned. As such, the early CNN-layer can be considered to be similar to the local contrast Weibull model that was previously noted to explain ERP amplitude early in time [27]. Higher in the visual hierarchy, the features represented in area V4 and IT become increasingly complex, containing shapes and object-like intermediate features [24]), which are thought to be processed later in time during feedforward visual processing. The complex features such as shapes, contours and even object-like features are captured by higher CNN layers [36].

Given the difference in complexities between CNN architectures, it is surprising that we observed no differences in maximal explained variance of brain responses by the individual layers of the CNNs. Potentially this can be attributed to equivalence of the layers across different CNN architectures that process the same extent of the image. For example, the receptive field size of layer 13 in CNN-15 is the same as layer 5 in CNN-7 [30]. However, increasing the number of layers improves the performance on object recognition for the ImageNet dataset [31], which is attributed to the additional computations (linear and non-linear) in deeper CNNs compared to shallower CNNs. Specifically, the additional non-linear computations increase the discriminative power of deeper CNNs. Consistently, we observed an improved correspondence between deeper CNNs and evoked brain responses when we evaluated the CNN architectures in their entirety.

Interestingly, this increased performance of the combined layers of CNN-15 in terms of explaining ERP responses as compared to the individual CNN layers was observed only later in time. Previous studies show that later in time (150 ms after visual input enters the retina), feedback and recurrent processing influence object-related processing [20]. While core object-related processing is hypothesized to be mediated by feed-forward processing [29], [33] our results therefore highlight the potential importance of recurrent processing in object recognition at later stages of the visual time-course. We speculate that the increased sensitivity might be explained by a recurrent feedback signal to lower brain areas that amplifies neural responses to fine-grained, lower-level information by grouping responses to object features and enhancing them in relation to other responses [37], [25], [20]. In sum, while previous results suggest a correspondence between feed-forward hierarchy of visual representations and CNNs, our results suggest that CNNs might also be suitable to investigate mechanisms of recurrent processing. Specifically, our results suggest that the gain in explanatory power with increased CNN depth is limited when it comes to explaining brain responses within the feed-forward visual sweep (i.e., before 150 ms), but that increased depth of CNN does help better explain neural activity later in time.

In conclusion, we show that 1) the temporal dynamics of representations in the human visual hierarchy are captured by convolutional neural networks (CNN) and 2) deeper CNNs contain expanded representations of low-to-intermediate features which adds substantial predictive power towards explain ERP responses later in visual processing. Going forward, a number of questions remain for CNNs as a model for information processing in the brain. The parameters of CNN models, such as number of layers, layer dimension, type of layers are chosen based on object recognition performance. How these parameters relate to the representations in the human visual system is poorly understood. While in the current study we investigated the number of layers, further work is required to understand the correspondence of the number of CNN layers to the human visual system. Additionally, CNN features themselves are computed in a feed-forward manner and feedback mechanisms are not present in standard CNNs. Clearly, human visual processing is not only rapid and dynamic, but also highly complex, with feedback and recurrent processing playing an important role in visual processing beyond feed-forward information extraction [14]. However, our results suggest that detailed research on CNN architecture and computations in relation to neuroimaging measurements may provide novel insights in the neural mechanisms underlying in visual object recognition in the human brain. Further research is required to disentangle the role of feed-forward and feedback processing in visual processing of objects in relation to the visual features represented by CNN models.

## Acknowledgments

This research is supported by COMMIT.

## SI Video 1

Layer 1 and 6 of CNN-7 correlation to ERP. We show the whole scalp visualization of 2 different layers from CNN-7 between 100 ms and 300 ms. Lower layer 1 is highly correlated in channel overlaying early visual cortex early in time, t = 110 to 120 ms while the higher layer 6 is correlated later in time, t = 160 to 170 ms in channels overlaying higher visual areas.

## Abstract published : VSS 2016

An abstract of the work here was presented at the Annual vision science meeting 2016.

## References

[1] Kenneth P Burnham and David R Anderson. Multimodel inference understanding aic and bic in model selection. Sociological methods & research, 33(2):261–304, 2004.

[2] Charles F Cadieu, Ha Hong, Daniel L K Yamins, Nicolas Pinto, Diego Ardila, Ethan A Solomon, Najib J Majaj, and James J DiCarlo. Deep neural networks rival the representation of primate it cortex for core visual object recognition. PLoS computational biology, 10:e1003963, 2014 Dec 2014.

[3] Cichy Aditya Khosla, Dimitrios Pantazis, Antonio Torralba, and Aude Oliva. Comparison of deep neural networks to spatio-temporal cortical dynamics of human visual object recognition reveals hierarchical correspondence. Scientific reports, 6, 2016.

[4] Radoslaw Cichy, Aditya Khosla, Dimitrios Pantazis, Antonio Torralba, and Aude Oliva. Mapping human visual representations in space and time by neural networks. Journal of vision, 15(12):376–376, 2015.

[5] Mark Everingham, Luc J. Van Gool, Christopher K. I. Williams, John M. Winn, and Andrew Zisserman. The pascal visual object classes (voc) challenge. International Journal of Computer Vision, 88(2):303–338, 2010.

[6] Sennay Ghebreab, Steven Scholte, Victor Lamme, and Arnold Smeulders. A biologically plausible model for rapid natural scene identification. In Advances in Neural Information Processing Systems, pages 629–637, 2009.

[7] Gabriele Gratton, Michael GH Coles, and Emanuel Donchin. A new method for off-line removal of ocular artifact. Electroencephalography and clinical neurophysiology, 55(4):468–484, 1983.

[8] Kalanit Grill-Spector, Tammar Kushnir, Talma Hendler, Shimon Edelman, Yacov Itzchak, Rafael Malach, et al. A sequence of object-processing stages revealed by fmri in the human occipital lobe. Human brain mapping, 6(4):316–328, 1998.

[9] Groen Sennay Ghebreab, Victor Lamme, and Steven Scholte. The role of weibull image statistics in rapid object detection in natural scenes. Journal of Vision, 10(7):992–992, 2010.

[10] Groen Sennay Ghebreab, Victor AF Lamme and H Steven Scholte. Spatially pooled contrast responses predict neural and perceptual similarity of naturalistic image categories. PLoS Comput Biol, 8(10):e1002726, 2012.

[11] Groen Sennay Ghebreab, Hielke Prins, Victor AF Lamme and H Steven Scholte. From image statistics to scene gist: evoked neural activity reveals transition from low-level natural image structure to scene category. Journal of Neuroscience, 33(48):18814–18824, 2013.

[12] Umut Güçluü and Marcel AJ van Gerven. Deep neural networks reveal a gradient in the complexity of neural representations across the ventral stream. The Journal of Neuroscience, 35(27):10005–10014, 2015.

[13] Kaiming He, Xiangyu Zhang, Shaoqing Ren, and Jian Sun. Delving deep into rectifiers: Surpassing human-level performance on imagenet classification. arXiv preprint arXiv:1502.01852, 2015.

[14] David J Heeger. Theory of cortical function. Proceedings of the National Academy of Sciences, 114(8):1773–1782, 2017.

[15] D. Hubel and T. N. Wiesel. Receptive fields, binocular interaction, and functional architecture in the cat’s visual cortex. Journal of Physiology, 160:106–154, 1962.

[16] Herve Jegou, Matthijs Douze, and Cordelia Schmid. Hamming embedding and weak geometric consistency for large scale image search. In Computer Vision-ECCV 2008, pages 304–317. Springer, 2008.

[17] Nancy Kanwisher and Etc Dilks. The functional organization of the ventral visual pathway in humans.

[18] Seyed-Mahdi Khaligh-Razavi and Nikolaus Kriegeskorte. Deep supervised, but not unsupervised, models may explain it cortical representation. PLoS Comput Biol, 10(11):e1003915, 11 2014.

[19] Alex Krizhevsky, Ilya Sutskever, and Geoffrey E. Hinton. Imagenet classification with deep convolutional neural networks. In Peter L. Bartlett, Fernando C. N. Pereira, Christopher J. C. Burges, Lon Bottou, and Kilian Q. Weinberger editors, NIPS, pages 1106–1114, 2012.

[20] Victor AF Lamme and Pieter R Roelfsema. The distinct modes of vision offered by feedforward and recurrent processing. Trends in neurosciences, 23(11):571–579, 2000.

[21] Adriana Olmos et al. A biologically inspired algorithm for the recovery of shading and reflectance images. Perception, 33(12):1463–1473, 2003.

[22] Andreas Opelt, Axel Pinz, Michael Fussenegger, and Peter Auer. Generic object recognition with boosting. Pattern Analysis and Machine Intelligence, IEEE Transactions on, 28(3):416–431, 2006.

[23] F Perrin J Pernier, O Bertrand, and JF Echallier. Spherical splines for scalp potential and current density mapping. Electroencephalography and clinical neurophysiology, 72(2):184–187, 1989.

[24] Kandan Ramakrishnan, H Steven Scholte, Iris I A Groen, Arnold W Smeulders, and Sennay Ghebreab. Visual dictionaries as intermediate features in the human brain. Frontiers in Computational Neuroscience, 8(168), 2015.

[25] Pieter R Roelfsema, Victor AF Lamme, and Henk Spekreijse. The implementation of visual routines. Vision research, 40(10):1385–1411, 2000.

[26] Olga Russakovsky, Jia Deng, Hao Su, Jonathan Krause, Sanjeev Satheesh, Sean Ma, Zhiheng Huang, Andrej Karpathy, Aditya Khosla, Michael Bernstein, et al. Imagenet large scale visual recognition challenge. International Journal of Computer Vision, pages 1–42, 2014.

[27] H Steven Scholte Sennay Ghebreab, Lourens Waldorp, Arnold WM Smeulders, and Victor AF Lamme. Brain responses strongly correlate with weibull image statistics when processing natural images. Journal of Vision, 9(4):29, 2009.

[28] Noor Seijdel, Kandan Ramakrishnan, Max Losch, and Steven Scholte. Overlap in performance of cnn’s, human behavior and eeg classification. Journal of Vision, 16(12):501–501, 2016.

[29] Thomas Serre, Aude Oliva, and Tomaso Poggio. A feedforward architecture accounts for rapid categorization. Proceedings of the national academy of sciences, 104(15):6424–6429, 2007.

[30] Karen Simonyan and Andrew Zisserman. Very deep convolutional networks for large-scale image recognition. arXiv preprint arXiv:1409.1556, 2014.

[31] Christian Szegedy, Wei Liu, Yangqing Jia, Pierre Sermanet, Scott Reed, Dragomir Anguelov, Dumitru Erhan, Vincent Vanhoucke, and Andrew Rabinovich. Going deeper with convolutions. CoRR, abs/1409.4842, 2014.

[32] Simon Thorpe, Denise Fize, and Catherine Marlot. Speed of processing in the human visual system. nature, 381(6582):520, 1996.

[33] Rufin VanRullen and Simon J Thorpe. Surfing a spike wave down the ventral stream. Vision research, 42(23):2593–2615, 2002.

[34] Dirk B Walther, Eamon Caddigan, Li Fei-Fei, and Diane M Beck. Natural scene categories revealed in distributed patterns of activity in the human brain. The Journal of Neuroscience, 29(34):10573–10581, 2009.

[35] Daniel L. K. Yamins, Ha Hong, Charles F. Cadieu, Ethan A. Solomon, Darren Seibert, and James J. DiCarlo. Performance-optimized hierarchical models predict neural responses in higher visual cortex. Proceedings of the National Academy of Sciences, 111(23):8619–8624, 2014.

[36] Matthew D Zeiler and Rob Fergus. Visualizing and understanding convolutional networks. In Computer Vision-ECCV 2014, pages 818–833. Springer, 2014.

[37] Karl Zipser, Victor AF Lamme, and Peter H Schiller. Contextual modulation in primary visual cortex. Journal of Neuroscience, 16(22):7376–7389, 1996.

